# Engineered Transcription Factor Binding Arrays for DNA-based Gene Expression Control in Mammalian Cells

**DOI:** 10.1101/2024.09.03.610999

**Authors:** A Zouein, B Lende-Dorn, KE Galloway, T Ellis, F Ceroni

## Abstract

Manipulating gene expression in mammalian cells is critical for cell engineering applications. Here we explore the potential of transcription factor (TF) recognition element arrays as DNA tools for modifying free TF levels in cells and thereby controlling gene expression. We first demonstrate proof-of-concept, showing that Tet TF-binding recognition element (RE) arrays of different lengths can tune gene expression and alter gene circuit performance in a predictable manner. We then open-up the approach to interface with any TF with a known binding site by developing a new method called Cloning Troublesome Repeats in Loops (CTRL) that can assemble plasmids with up to 256 repeats of any RE sequence. Transfection of RE array plasmids assembled by CTRL into mammalian cells show potential to modify host cell gene regulation at longer array sizes by sequestration of the TF of interest. RE array plasmids built using CTRL were demonstrated to target both synthetic and native mammalian TFs, illustrating the ability to use these tools to modulate genetic circuits and instruct cell fate. Together this work advances our ability to assemble repetitive DNA arrays and showcases the use of TF-binding RE arrays as a method for manipulating mammalian gene expression, thus expanding the possibilities for mammalian cell engineering.

## Introduction

The ability to precisely control gene expression is key to many biotechnology and synthetic biology applications. Precise and dynamic control over gene expression can be achieved using transcription factors (TFs) which modulate gene expression levels by recognizing and interacting with specific DNA recognition elements (REs) near promoters, playing a major role in cellular maintenance, adaptation to environmental changes, and cell fate transitions (Lambert et al., 2018). As a result, modulating TFs is a potent means to rewire endogenous gene expression programs for cellular reprogramming or investigating cell states (Bhatt et al., 2023; Joung et al., 2023; Vogel et al., 2019; N. B. Wang et al., 2020).

TF engineering is often based on the overexpression of synthetic (Bhatt et al., 2023; Donahue et al., 2020; Fidanza et al., 2017; Krawczyk et al., 2020) or native master TF genes (Joung et al., 2023; Takahashi & Yamanaka, 2006). However, this strategy often raises safety concerns and is prone to gene silencing or oncology (Bhatt et al., 2024; Bolognesi & Lehner, 2018). Moreover, as intracellular resource competition becomes more critical for the success of many applications (Qin et al, 2023; Di Blasi et al., 2023; Eisenhut et al., 2024; R. Zhang et al., 2021), tools that do not rely on intracellular gene expression resources for action may have advantages.

In this context, transcription factor binding site ‘decoys’ offer a promising alternative. This group of synthetic DNA constructs are designed to contain REs that competitively sequester TFs away from their target recognition sites in the host genome. These decoys are also sometimes known as ‘sponges’ and can be either (i) self-replicating plasmids, (ii) decoy oligonucleotides or (iii) RE array plasmids (Lee & Maheshri, 2012; Rad et al., 2015; Wan et al., 2020a; X.-F. Wang & Calame, 1986). Self-replicating plasmids use viral origin of replications to achieve high RE numbers in cells but have a low number of sites per plasmid (X.-F. Wang & Calame, 1986). Short oligonucleotides also contain a low number of sites per molecule but are transiently supplied to cells at high concentrations (Rad et al., 2015). The final class of decoy, RE arrays, increase RE number per molecule by concatenating REs into arrays to achieve a high local concentration of TF binding to each plasmid.

Both the self-replicating plasmid and short oligonucleotide types of decoys have previously been used in mammalian cell research (Crinelli et al., 2004; Rad et al., 2015; X.-F. Wang & Calame, 1986) but face challenges that limit their broad adoption. Decoy oligonucleotides often lead to immunogenicity and poor intracellular stability (Rad et al., 2015), making them ineffective for some applications. Self-replicating plasmids are longer-lived but depend on viral components, which complicates their use in-vivo (Park et al., 1999; X.-F. Wang & Calame, 1986). Notably, neither self-replicating plasmids nor oligonucleotides make use of TF cooperative binding, a key feature of eukaryotic TFs. TFs have improved affinity to DNA regions containing multiple sequential REs (Bashor et al., 2019; Bragdon et al., 2023; Ibarra et al., 2020; MacCarthy et al., 2024), and so repeats of REs should improve TF sequestration.

In prior work in yeast and bacteria models, the use of arrays of repeated REs for specific TFs has been used to achieve precise control of gene expression, enhancing performance through reduced expression leakage, increased output, and improved dynamic range (Lee & Maheshri, 2012; Wan et al., 2020). However, despite the successes in microbial systems, the use of similar TF RE arrays in mammalian cells has not yet been demonstrated. To some degree, this is due to the unique challenges of assembling plasmids with many repeats of a single DNA sequence. Constructing DNA with repetitive sequences is technically challenging due to various factors, including DNA recombination and sequence instability. These issues often occur in cloning hosts like *E. coli* and when methods such as PCR are adopted.

Here we establish the use of repetitive TF RE array plasmids as a tool for tuning gene expression in mammalian cells, first demonstrating this with TetR-based gene circuits. We then overcome current limitations by developing an assembly strategy that can be used to construct plasmids containing up to 256 REs. We demonstrate multiple applications using RE arrays as decoys for targeted TFs, enhancing our comprehension of TF functionality. Moreover, we observe the influence of these arrays on the tuning of synthetic gene circuits and their potential to guide cells towards specific states.

## Results & Discussion

### TF RE array plasmids can sequester synthetic TFs in mammalian cells

To demonstrate the concept of TF RE arrays for tuning gene regulation in mammalian cells, we first evaluated the impact of TetO-array containing plasmids on synthetic gene circuits that use TetR-based TFs. We chose the TetR group of TFs as they are well studied and RE arrays that bind TetR TFs have been constructed, tested and shown to tune genetic circuits in bacteria and yeast (Lee & Maheshri, 2012; Wan et al., 2020).

To assess the functionality of the RE arrays, we use a plasmid encoding a gene circuit that constitutively expresses the synthetic TF, rtTA (reverse tetracycline transactivator), using this to trigger expression of a fluorescent reporter (EGFP) by binding TetO sites in the upstream region of the pTet-A promoter in the presence of doxycycline (dox) (**Figure 1a**). This is co-transfected along with a competitor plasmid to identify designs capable of deterring EGFP expression detected by flow cytometry 48 hours post-transfection. Excess TetO sites provided by a competitor plasmid would presumably titrate rtTA away from pTet-A, reducing its ability to active EGFP expression, whereas plasmids not containing TetO sites, a plasmid only containing regions for plasmid propagation in bacteria for example (pEmpty) should not impact the expression of EGFP (**Figure 1a**). Competitor plasmids with an increasing number of TetO repeats in series showed a reduction in EGFP output compared to the pEmpty conditions, which lacked TetO sites in HEK-293T cells (**Figure 1b**) and CHO-K1 cells (**Supplementary Figure 1a**) as measured by the Relative Fluorescence Units (RFU) of the singlet population.

**Figure 1:**
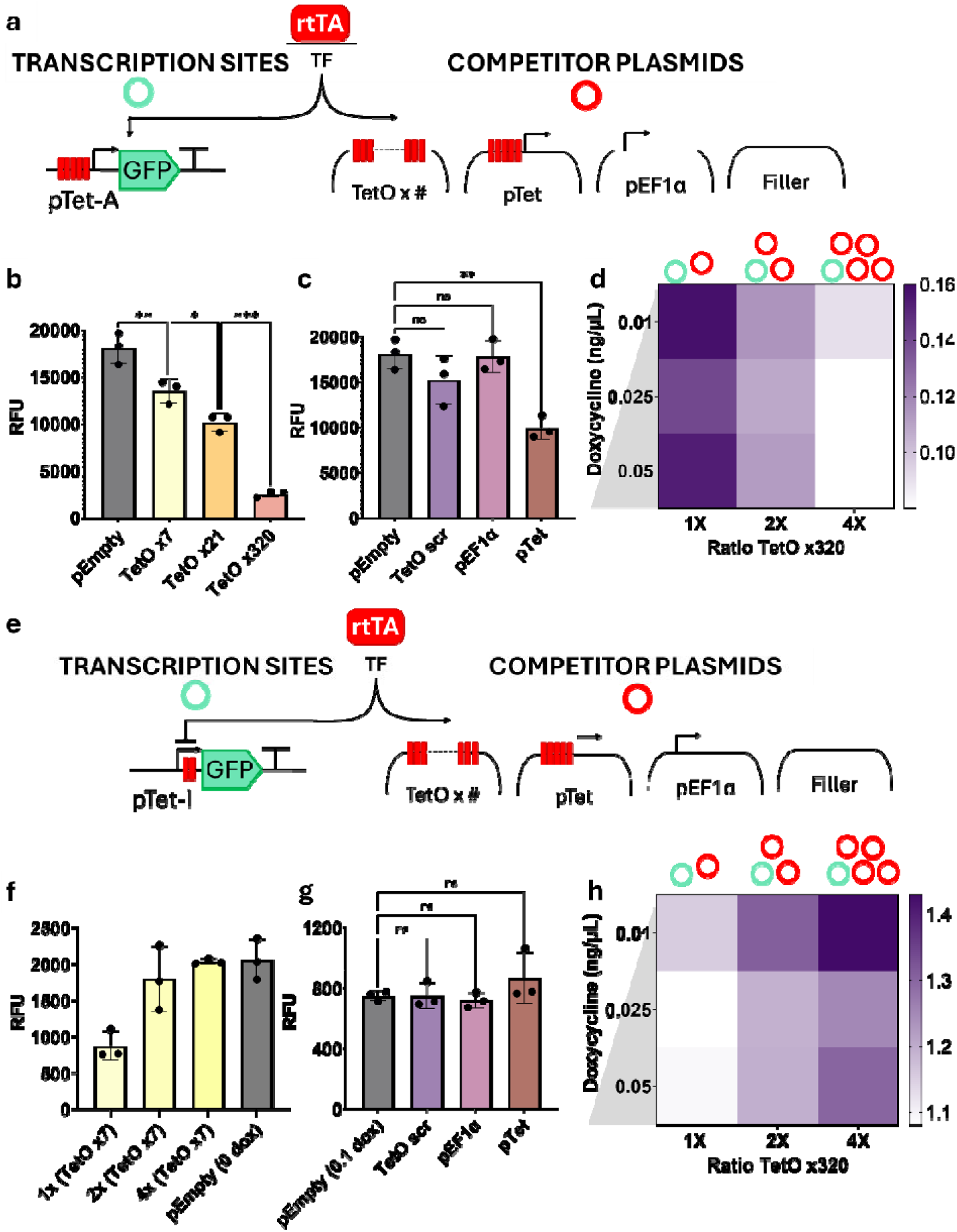
Effect of TetO RE arrays on synthetic circuits in HEK-293T cells. **a.** In the presence of doxycycline, a finite pool of rtTA can bind to either the competitor plasmids or to the gene of interest at promoter, pTet-A, where it activates transcription of EGFP. **b.** EGFP expression in Relative Fluorescent Units (RFU) measured from cells in 0.05 ng/µL of dox after transfection with equimolar amounts of competitor plasmids containing 0 (pEmpty), 7, 21 or 320 TetO sites. **c.** EGFP expression measured from cells in 0.05 ng/µL of dox after transfection with equimolar amounts of control plasmids pEmpty, TetOscr, pEF1a or pTet. **d.** Heatmap depicting fold change relative to pEmpty for various circuit-to-array plasmid DNA ratios across different dox concentrations. **e.** In the presence of dox, a finite pool of rtTA can bind to either the competitor plasmids or to the gene of interest promoter, pTet-I containing 2 TetO sites within the core promoter, repressing expression of EGFP. **f.** EGFP expression measured from cells in 0.05 ng/µL of dox after transfection with equimolar, double or quadruple competitor plasmid containing 7 TetO sites and 0.05 ng/µL dox as compared to pEmpty in absence of dox (i.e. showing maximum expression). **g.** EGFP expression measured from cells in 0.05 ng/µL of dox after transfection with controls TetOscr, pEF1a or pTet as compared to pEmpty in 0.1 ng/µL dox (representing maximum repression). **h.** Heatmap depicting the fold change of expression relative to the pEmpty control when increasing ratio of TF RE array relative to active transcription sites are added for a range of dox concentrations. Data represent mean values from three independent biological replicates while error bars represent ± standard deviation.

In HEK co-transfecting equimolar amounts of the circuit along with plasmid containing seven sequential TetO sites (TetOx7) led to a 30% decrease in reporter expression. A more substantial reduction of 53% was observed with the TetOx21 array. The most pronounced effect was seen with the TetOx320 plasmid, sourced from Wan et al, which decreased expression to just 9% of the levels seen in the absence of the TetO sites (pEmpty). A similar trend was observed in CHO-K1 cells, where TetO array plasmids repressed the circuit by 23% with TetOx7 or up to 54% when TetOx320 array is used (**Supplementary Figure 1a**) indicating that the trends observed are not simply due to a single specific cell-line background.

As further controls, we showed that two other plasmid designs also behave like pEmpty with no significant effect on the circuit (**Figure 1c**); (1) pEF1α, a plasmid encoding a strong constitutive mammalian promoter that tests whether increased docking of basic transcription machinery at promoter elements accounts for changes in EGFP expression, and (2) TetOscr, a plasmid with 48 consecutive ‘scrambled’ TetO sites where each site has the same nucleotide content as TetO but in a different order. This latter control verifies that any observed changes in EGFP expression are strictly due to site-specific binding of rtTA to its target binding sites. Finally, we also tested whether a plasmid encoding the full pTet-A promoter with its seven TetO sites had an effect on EGFP expression. The presence of this plasmid did indeed lead to a decrease in EGFP, but the measured decrease was equivalent to that of the 7xTetO array suggesting that the effect was entirely due to TetO sites (**Figure 1c**).

To further explore whether varying the transfected amounts of plasmid DNA could also tune circuit output, we added 1x, 2x, or 4x concentrations of the TetOx320 RE array and observed the effects on the circuit (**Figure 1d, Supplementary Figure 1b, Supplementary Figure 1c**). The results in **Figure 1d** indicate that increasing the amount of TetOx320 RE array plasmid in HEK-293T led to a decrease in fluorescence detected from the circuit. This held true for a range of doxycycline concentrations, representing an increased pool of TF capable of sequestration by TF RE array. Similarly, in CHO-K1 cells the addition of more competitor plasmid acted as a method to tune the dynamic range of the circuit, with adding quadruple the molar amount of RE array plasmid having the biggest impact on EGFP expression (**Supplementary Figure 1c**). Interestingly, RE array plasmid did not reduce the level of basal expression of EGFP in the absence of dox, suggesting leaky expression from pTet-A is not due to rtTA binding in the absence of its inducer (**Supplementary Figure 1b, c**).

Having confirmed that TetO array plasmids can modulate simple transcriptional activation circuits, we next sought to understand the broader applicability of RE arrays on circuits where rtTA represses mKate reporter protein expression. Here, the pTet-I promoter was used, containing two TetO sites in its core region which when bound by rtTA acts as a steric hindrance blocking transcription initiation (Yao et al., 1998) (**Figure 1e**). In the absence of dox, rtTA cannot bind and reporter resulting in higher mKate expression (2000 RFU), whereas this decreased to 750 RFU when 0.1 ng/µL of dox is added (**Figure 1f,g**). The addition of TF RE array plasmids should sequester rtTA/dox complexes and reduce repression, so that mKate expression increases. Indeed, increased expression in the presence of RE array plasmids was observed, both when longer arrays were used and when more plasmid DNA was transfected (**Figure 1f,h**). Transfection of control plasmids did not affect the gene circuit (**Figure 1g**), indicating differential expression is due to TF/RE interactions.

The effects of TetO arrays on TetR-based gene circuits with positive autoregulation (Di Blasi et al., 2023) and negative autoregulation (Szenk et al., 2020) were also tested and showed the expected effects (**Supplementary Figure 2a-c**). Transfection of the positive autoregulation circuit (Di Blasi et al., 2023), in which rtTA expression is controlled by the pTet-A promoter, required only a small amount of TetO array plasmid (0.5x the equimolar amount of the circuit plasmid), to modulate expression. This indicates the sensitivity of this circuit to perturbation by RE array plasmids (**Supplementary Figure 2a**). In contrast, the TetR negative autoregulation circuit required four times more RE arrays containing 320 TetO sites to see less than 25% increase in expression (**Supplementary Figure 2c**). Using such a high amount of RE array plasmid in this experiment came at the expense of cell viability (**Supplementary Figure 2c**). As this concentration of the array was used in **Figure 1** and did not lead to cell death, cytotoxic protein overexpression by perturbing negative autoregulation as reported in (Chaturvedi et al., 2022) may be leading to this.

Lastly, we extended our work with TetO array plasmids to show that they can act with other synthetic TFs binding them. For this we used dCas9-VPR (Fidanza et al., 2017), targeting this activating TF to TetO sites via expression of a specific guide RNA. As expected, the presence of RE array plasmids reduced the ability of the dCas9-VPR to activate gene circuit expression (**Supplementary Figure 2b**). Altogether we showed that a variety of different gene regulatory circuits can be predictably perturbed by synthetic RE arrays in mammalian cells and their action is due to the REs and their relative numbers and concentrations.

### Assembly of arrays of repetitive sequences by CTRL

To extend beyond TetO arrays and facilitate the use of RE array plasmids that can sequester any TF of choice including mammalian ones, we developed a standardized assembly workflow for constructing plasmids with repetitive synthetic arrays. Given the complexities associated with assembling repetitive DNA sequences, as highlighted by toolkits summarized in Zouein et al., (2021) we saw an opportunity to develop an approach to overcome these limitations and expedite the construction of repetitive plasmids for broader use.

CTRL (**C**loning **T**roublesome **R**epeats in **L**oops) is a cloning workflow designed with the following features to improve the modular assembly of repetitive sequences. Firstly, the multiplication of repetitive sequences is achieved in cycles, allowing the controlled assembly of specific numbers of repeats. Secondly, CTRL is backward compatible with the MTK (**M**ammalian **T**ool**k**it, Fonseca et al., 2020) a widely available Golden Gate (GG) toolkit with a library of DNA parts that can be assembled in a modular fashion to construct mammalian gene circuits. This allows the addition of RE arrays as modular parts that can be added into more complex gene circuit designs. It also facilitates potential integration of RE arrays into mammalian genomes, as the MTK system uses plasmid backbones that enable direct transfer into integration vectors.

To address the challenges associated with handling repetitive sequences, which are susceptible to deleterious mutation and recombination, CTRL plasmid backbones were chosen to have a low copy number in *E. coli* to minimize risk of replication slippage (Richards, 2013). A series of neutral DNA spacers were also incorporated between REs to further reduce the likelihood of deleterious homologous recombination during plasmid propagation and amplification in bacterial hosts (Wan et al., 2020b). An additional feature, as seen in other Golden Gate toolkits (J. P. Fonseca et al., 2019; Martella et al., 2017), is the use of a drop-out cassette to facilitate the identification of successful cloning events, in this case one constitutively expressing a red fluorescent protein in *E. coli*. The drop-out cassette allows users to visually differentiate between white colonies, in which DNA has assembled into the plasmid, and red bacterial colonies which are the original vector plasmid having not taken in new DNA. Finally, the use of Golden Gate assembly further allows the user to customize CTRL for applications that require the absence of cloning scars in the final assembled DNA.

The CTRL workflow begins with the custom DNA synthesis of a compatible insert (grey box, **Figure 2a**) containing TF REs separated by neutral spacer sequences. RE sequences and spacers used in this study can be found in **Supplementary Tables 1 and 2**, respectively. Synthesized inserts are then assembled into ampicillin resistant entry vectors via internal BsmbI cut sites (**Supplementary Table 3**) and cloned via standard Golden Gate (GG) assembly methods. Repeating the process in subsequent ‘loops’ multiplies the number of RE repeats in the plasmid. RE repeats can be doubled by using pairwise-assembly (via the robust BioBrick assembly method) or can be quadrupled in each cloning loop using GG assembly (**Figure 2a**). Cloning loops are repeated until the desired number of REs in an array is achieved. In each round, successful assemblies can be identified by the loss of red fluorescence in *E. coli* colonies followed by plasmid extraction from a sample of white colonies, and standard gel electrophoresis analysis to check the RE array size. Final RE array assemblies can then be further modified to suit final applications by substituting the CTRL cloning vector backbones for plasmid backbones of interest from the MTK system (Addgene Kit #1000000180).

**Figure 2:**
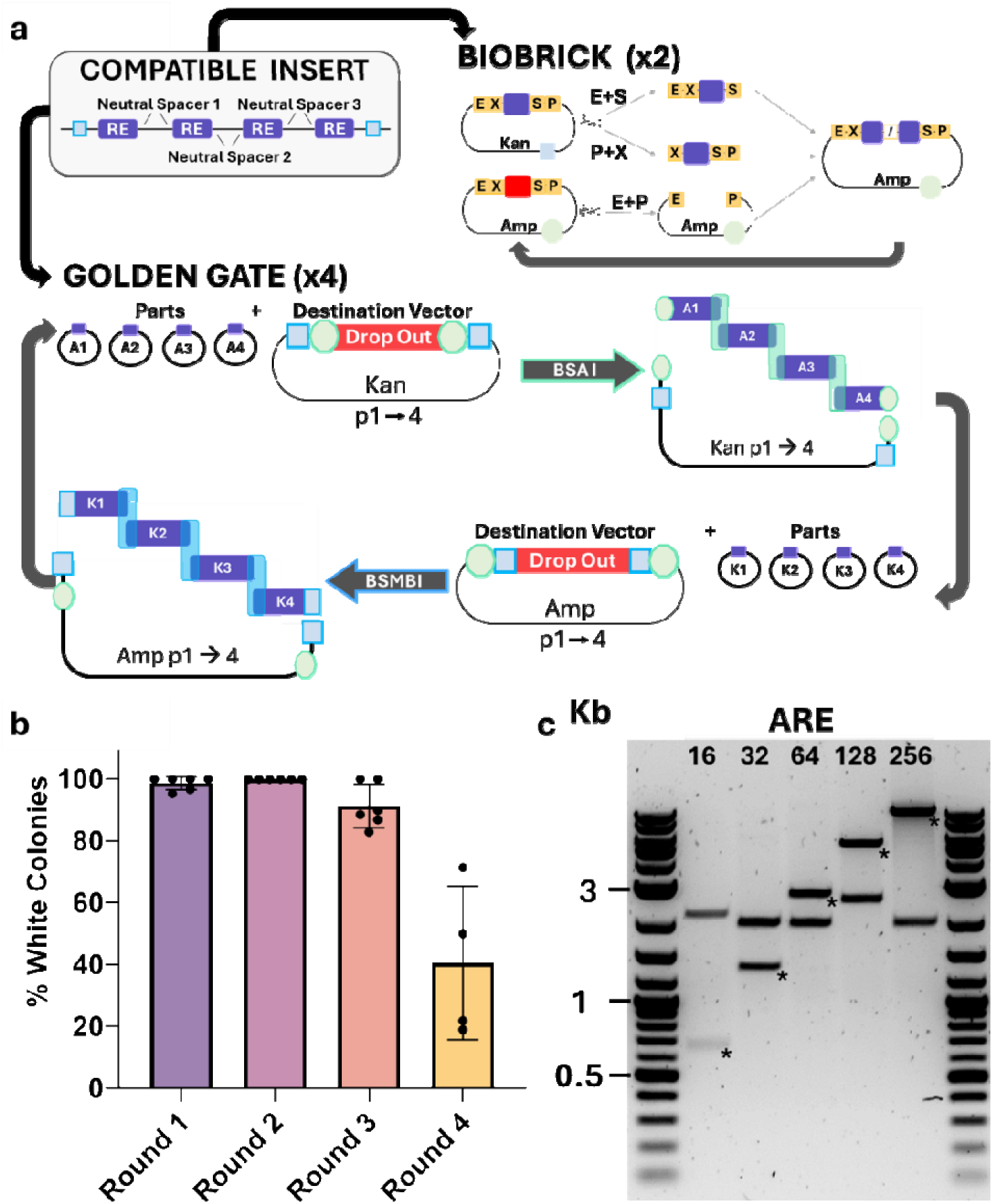
Development of CTRL for assembly of repetitive DNA sequences. **a.** Schematic of CTRL workflow. Compatible inserts containing TF REs, neutral spacers and restriction sites are used for domestication into CTRL which can multiply inserts using BioBrick function or the Golden Gate function. BioBrick: A portion of insert is digested with EcoRI (E) and SpeI (S) while another portion is digested with PstI (P) and XbaI (X). A CTRL backbone containing opposite resistance cassette (Amp) is digested with EcoRI and PstI. Ligation of these 3 parts yields a plasmid containing double the number of repeats separated by an uncleavable scar, annotated as /, from the ligation product of XbaI and SpeI overhangs. Golden Gate: BsaI reactions cut inserts from Ampicillin positions 1 through 4 and assemble into receivers with Kanamycin resistance and GG reactions using BsmBI to cut inserts from Kanamycin positions 1 through 4 and assemble into receivers with ampicillin resistance quadrupling RE array. **b.** Percent white colonies for sequential cycles of cloning as an indicator of the assembly’s rate of success. **c.** Agarose gel image of EcoRI and PstI restriction digested repeat containing CTRL products for mammalian RE ARE. Repeat-containing fragments are marked with an asterisk (*), while other bands represent either ampicillin or kanamycin resistance containing CTRL backbones. Lanes 1 and 7 contain 1 Kb Plus DNA ladder (NEB) as a molecular weight marker. Data represent mean values from at least four independent biological replicates while error bars represent ± standard deviation.

The GG assembly route of CTRL can multiply repeats by four in each cycle (**Figure *2*a**) and requires alternating the use of vectors with different antibiotic selection markers to ensure successful assembly. Initially, the inserts are introduced into four position vectors containing ampicillin resistance, labeled as positions a1 through a4 representing the ligation order of the inserts. In the GG reaction these plasmids are then combined with a destination vector containing kanamycin resistance and BsaI enzyme to ligate position a1 to position a2 to position a3 to position a4 to the destination vector. To create all the necessary parts for the next assembly, this reaction is set up for four destination vectors (k1,k2,k3,k4), that generate the ligation order in the next GG reaction. The plasmids assembled from this process now serve as the parts for the subsequent round of GG cloning which ligates position k1 to position k2 to position k3 to position k4 to a destination vector containing ampicillin, of which there are four. Therefore, assembly into ampicillin positions creates the necessary context for the next round of cloning into kanamycin resistance vectors and vice versa. To proceed with the next round of GG cloning, assembly into all four receiver vectors must be successful to form the parts required for the subsequent assembly.

The standardized BioBrick approach, integrated into CTRL, facilitates shorter pairwise assemblies using XbaI and SpeI isocaudamers, which produce compatible cohesive ends. This forms plasmids with an intermediate length of repeat units separated by a scar (denoted as / in **Figure *2*a**) and reforms all required restriction sites for subsequent rounds of cloning (Shetty et al., 2008; Sladitschek & Neveu, 2015).

To assess the efficiency of our CTRL workflow and its robustness to variations in DNA sequence and length, we demonstrated the assembly of RE array plasmids for three different mammalian TFs, namely NRF2, Prrx1 and Tead1. These were selected due to their reported significance in the regulation of mammalian cell response to oxidative stress and direct conversion, respectively (Cahan et al., 2014; R. Wang & Perez, 2020). RE sequences used for NRF2, Prrx1 and Tead1 are reported in Supplementary Table 1. After four rounds of CTRL assembly, arrays of 256 repeats for each RE were successfully constructed. The first, second, and third round of assembly yielded correct arrays over 92% of the time, whereas only 40% of colonies contained potentially correct arrays in the fourth round of cloning (**Figure 2b**), as calculated from counting white vs. red colonies on agar plates.

In our experiments we stopped cloning cycles after four CTRL rounds had generated arrays with 256 RE, however, we have no reason to believe that this is the limit of CTRL, with longer arrays likely being possible with further rounds. Verification of assemblies was performed by restriction digestion of plasmid DNA extracted from bacterial colonies (**Figure 2c, Supplementary Figure 3**) and DNA sequencing (data not shown, Full Circle Labs). Through CTRL cloning using both Biobrick and Golden Gate modalities, plasmids containing 16, 32, 64, 128 and 256 sequential ARE sites (Antioxidant Response Element) for targeting of the mammalian TF NRF2 were built. The final set of ARE array plasmids built are shown in **Figure 2c**, digested with restriction enzymes flanking the arrays, where the band containing the repeat array is indicated by asterisks. The size match of these bands with simulated assemblies confirmed the successful assembly of the desired DNA fragments. Similar verification was performed on Prrx1 and Tead1 REs (**Supplementary Figure 3**). Verification of assemblies was performed by plasmid extraction from colonies, followed by restriction digestion and gel electrophoresis analysis (**Figure 2c**, **Supplementary Figure 3**) and whole plasmid DNA sequencing.

For NRF2-sequestering RE arrays, we used CTRL via both the BioBrick and Golden Gate methods to construct plasmids containing 16, 32, 64, 128 and 256 sequential ARE (Antioxidant Response Element) sites to which the mammalian NRF2 TF binds. The gel electrophoresis verification of the final set of ARE array plasmids built is shown in **Figure 2c**, digesting with restriction enzymes that flank the arrays. The size match of these bands with the sizes given from simulated assemblies confirmed the successful construction of the desired RE arrays. Similar verification was performed on RE array plasmids constructed for the Prrx1 and Tead1 TF binding sites (see **Supplementary Figure 3**).

Building on our success of building RE arrays we set to test CTRL features that facilitate using RE arrays as parts of more complex circuits for mammalian applications. As Prrx1 and Tead1 support fibroblast identity, recruitment of these TFs to RE arrays may influence direct conversion by sequestering these TFs from their native targets (Cahan et al, 2014). To examine this hypothesis, we encoded the RE arrays into a lentiviral backbone, along with an additional transcriptional unit expressing the reporter mRuby2. The decoy arrays for Prrx1 or Tead1 were delivered to primary Hb9::GFP transgenic mouse embryonic fibroblasts (MEFs). We confirmed these sequences were delivered and integrated into the genome by their mRuby2 fluorescence (**Supplementary Figure 4a-d**).

To examine the influence on direct conversion, we quantified the number of cells that activated the Hb9::GFP motor neuron reporter at 14 days post infection (dpi). Addition of motor neuron transcription factors drives cells to activate the Hb9::GFP reporter as the cells adopt the motor neuron identity (Son et al.,2011; Babos et al.,2019; Wang et al,.2023). Examining cells that expressed mRuby2, the delivery of the RE arrays did not change the fraction of cells that converted to motor neurons (**Supplementary Figure 4e-g**). The system remained robust against the influence of Prrx1 and Tead1 RE arrays, suggesting that recruitment to these randomly integrated sites does not affect the ability of Prxx1 and Tead1 to support fibroblast identity. Putatively, these data suggest that synthetic promoters composed of these arrays may report on TF levels without destabilizing fibroblast identity (Johansson et al., 2017; Wu et al., 2019).

### ARE array plasmids sequester NRF2 in mammalian cells

After confirming that CTRL can construct RE arrays with variations in size, DNA sequence, RE length and content, we next sought to test the impact of an array designed to interact with endogenous gene regulation in mammalian cells.

The native transcription factor NRF2 binds to sites in the genome known as AREs and alters gene expression from regions with these sites under oxidative stress conditions (Ganner et al., 2020). In non-oxidative conditions, NRF2 is sequestered in the cytoplasm and actively degraded. Here, we used arrays with 12, 16, 32, 64 and 128 copies of an ARE sequence to test the effect of NRF2 sequestration when cells are under oxidative stress, which we induced by addition of Auranofin, a potent NRF2 inducer (Johansson et al., 2017). Addition of RE arrays containing ARE sites should inhibit the action of NRF2 by reducing its binding to its genomic locations under conditions of oxidative stress. This would be expected to hamper the ability of a cell to survive the stress environment leading to increased cell death (Liu et al., 2023; J. Zhang et al., 2017) (**Figure 3a**).

**Figure 3:**
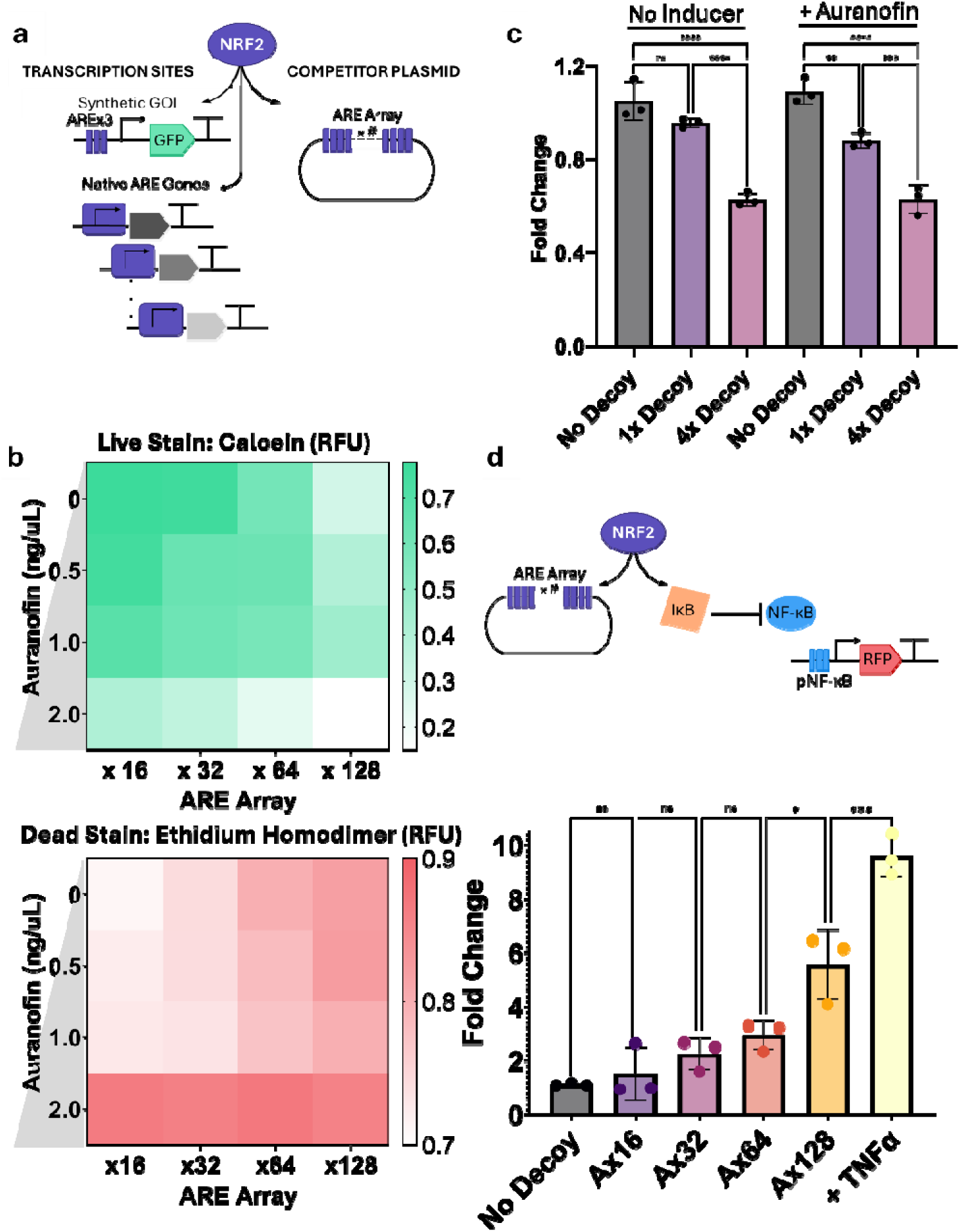
Use of ARE array plasmids to induce cell death. **a.** Schematic representation of NRF2 binding to active transcriptional regions or ARE array plasmids containing Antioxidant Response Elements (AREs). **b.** Heatmaps representing the viability of cells in response to varying lengths of ARE arrays and auranofin inducer. Results of calcein live staining (heatmap on the left) ranging from green to white, indicate the proportion of live cells where a darker shade of green signifies a higher proportion of live cells in culture 48hr post-transfection. Results of ethidium homodimer-1 (heatmap on the right) ranging from red to white, indicate the proportion of dead cells in culture where darker shades of red indicate a larger proportion of dead cells. **c.** Fold change in GFP expression from cellular NRF2 sensor in the presence or absence of 0.5 µM Auranofin comparing effect in the absence of AREs (black bars) 1x AREx12 array (purple bars) or 4x AREx12 array (pink bars). Fold change is calculated by the ratio of GFP expression levels to that with no AREs, no inducer condition. A fold change greater than 1 indicates upregulation, while a fold change less than 1 indicates downregulation of GFP. **d.** Schematic of NRF2 crosstalk with the TF NF-κB through its inhibitor IκB. NF-κB can be detected by expression of reporter from intracellular sensor. The bar graph displays the fold change in expression from NF-κB controlled reporter in the presence of ARE arrays or 20 ng/mL TNFα positive control compared to no RE condition. Data represent mean values from at least three independent biological replicates while error bars represent ± standard deviation.

Building on our observations sequestering rtTA with RE arrays, we first explored whether changing the length of ARE arrays would proportionally tune the amount of NRF2 available for genomic binding. We transiently transfected RE arrays containing 16, 32, 64 or 128 AREs and detected the effect of NRF2 sequestration on HEK-293T viability 48 hours post-transfection in the presence of 0, 0.5, 1 or 2 µM Auranofin. Cell viability in the presence of an RE array was compared to the control, pEmpty, and was assessed using commercial live/dead staining (**Figure 3b**) using calcein to stain live cells (green heatmap) and Ethidium Homodimer 1 to stain dead cells (red heatmap). When analysing our data, we observed that smaller RE arrays affected cell death at higher concentrations of Auranofin, whereas larger RE arrays more consistently led to cell death across all Auranofin concentrations tested (**Figure 3b**). Notably, the addition of AREx128 plasmid resulted in death of >80% of cells even in the absence of Auranofin. Based on this observation, we did not attempt testing of longer RE arrays than 128 repeat arrays.

We then sought to determine whether, similar to our study with TetR TFs, the addition of different ratios of smaller ARE arrays could also sufficiently influence NRF2 regulation. To easily track the activity of intranuclear NRF2 levels, we used a previously developed NRF2 responsive promoter (Johansson et al., 2017) that directs expression of EGFP (**Figure 3a**). Transfection with a 4:1 ratio of AREx12 plasmid to NRF2 sensor plasmid led to 40% reduced expression from the responsive promoter both in the presence and absence of 0.5 µM Auranofin. In contrast, no significant difference in expression was seen when equal amounts of ARE array and sensor plasmids were transfected (**Figure 3c**). This observation deviated from the trend seen with TetR sequestration (**Figure 1**), where equimolar amounts of short arrays altered the expression from the reporter. Potentially this could be due to the different binding characteristics of the tight-binding prokaryotic TF (TetR) compared to a mammalian TF. The latter class more typically being low-affinity bindings that achieve high specificity by cooperatively binding to multiple RE sites (Andreani et al., 2024).

We next devised an experiment to observe how RE array control of NRF2 levels can be used to perturb a native gene circuit. For this we characterised the change in the widely studied TF, NF-κB which is known to be regulated by NRF2 (Gao et al., 2022) (Figure 3d). NRF2 inhibits the expression of NF-κB by inducing the expression of its inhibitor (Gao et al., 2022; Hohmann et al., 2020; R. Wang & Perez, 2020). And so, in principle, suppression of NRF2, by ARE array plasmids, should allow more NF-κB to accumulate in the nucleus where it acts as a TF, binding cognate genomic regions. To test this, we designed an NF-κB sensor similar to previously reported promoter (Johansson et al., 2017), where NF-κB triggers the expression of a synthetic transcriptional unit coding for the expression of 3x mCherry (Johansson et al., 2017). As a positive control test for this construct, we also tested in presence of TNFα, a known inducer of NF-κB (**Figure 3d**, yellow bar). As expected, addition of TNFα increased expression from the sensor construct by over 8-fold. Meanwhile, addition of the ARE array plasmids increased the signal detected from the sensor in a dose-dependent manner. More specifically, AREx16 and AREx32 plasmids gave a 2-fold increase, while AREx64 led to an increase just under 3-fold. Notably, the AREx128 TF-array plasmid significantly enhanced reporter expression by more than 5-fold compared to control conditions with no RE array. Our data suggest that increasing the length of the array results in a progressive enhancement in the signal.

## Discussion

Synthetic control of gene regulation in mammalian cells typically relies on genetic constructs that require the expression of proteins like TFs for their function. While this standard approach to synthetic gene control is broadly adopted, the burden caused from additional gene expression by the host can be overlooked (Di Blasi et al., 2023; Van Craenenbroeck et al., 2000). Here we developed an alternative approach using plasmids containing repetitive arrays of TF REs that competitively sequester TFs of interest from synthetic or genomic RE sites. These TF RE arrays do not rely on host gene expression resources to function, are diluted by cell division, and are not amplified intracellularly, which means the quantity of DNA delivered is the effective dose.

We found that RE arrays against commonly used synthetic TFs, dCas9 and Tet TFs, are effective in achieving differential expression of reporters, by competitively binding TFs in mammalian cell lines. Furthermore, TetO array plasmids altered the expression of GOIs with different regulatory topologies, including those with autoregulatory feedback. These results highlighted the different impact of RE arrays on TFs with distinct autoregulatory mechanisms. Specifically, we observed that co-transfecting TF RE arrays with negative autoregulatory circuits resulted in cell death, indicating a robust but lethal resistance to perturbation. In contrast, positive autoregulatory circuits were more susceptible to expression perturbation by TF RE arrays, demonstrating a higher sensitivity to regulatory interference. These results may represent the expected outcome of using RE arrays on similar native regulatory topologies. For example, using RE arrays against a native TF regulated by negative autoregulation may lead to open-loop lethality whereas using RE arrays against TFs regulated by positive autoregulation may be especially susceptible to expression perturbation.

To widen the use of RE array tools, we then developed CTRL, a method for assembling long repetitive sequences. While advances in DNA synthesis have offered an alternative to traditional molecular cloning techniques, synthesis of repetitive sequences remains a challenge (Hoose et al., 2023). Some mammalian DNA assembly toolkits can handle repetitive sequences (Chu et al., 2011; Gogolok et al., 2016; Noskov et al., 2011; Sladitschek & Neveu, 2015) but either take time to achieve long arrays, rely on error-prone enzymes or do not generate specific numbers of repeats. CTRL is an iterative assembly method designed for handling of DNA repeats and can be done either by 4-fold expansion using GG assembly or 2-fold using the more simple-BioBrick method. While BioBrick is slower to achieve large RE arrays it has been shown previously to be a robust way to ‘chain’ constructs together into larger sizes (Sladitschek & Neveu, 2015). In implementing CTRL, we assembled RE array plasmids for several mammalian TFs, specifically NRF2, Prrx1, and Tead1. The produced plasmids varied in sequence and length, demonstrating the robustness and efficiency of CTRL in building arrays containing up to 256 REs. CTRL facilitates use of RE arrays in various applications as arrays can be combined with other circuit components to achieve goals. However, given the repetitive nature of RE arrays, one must consider how additional components may interact with the repeats. For example, viral vectors which are prone to recombination may destabilize RE arrays (Al-Allaf et al., 2013).

We faced this challenge when incorporating Prrx1 and Tead1 arrays within lentiviral vectors for delivery to MEF cells. Given that Prrx1 and Tead1 are known to support fibroblast identity, we were keen to verify the feasibility of lentiviral integration within MEF genome and the impact on these cells. While we could clone plasmids bearing 16 and 32 RE arrays for Prrx1 and 12 and 24 RE arrays for Tead1, we struggled to assemble plasmids with a higher number of RE repeats within lentiviral vectors. While smaller arrays were successfully delivered to the MEF genome, they did not affect cellular identity, highlighting the need to assess the impact of higher repeat numbers, different RE sequences or higher multiplicity of infection.

Finally, we used CTRL-assembled ARE array plasmids to alter the regulatory action of the native TF NRF2 to inhibit the protective response to oxidative conditions and to change how it interacts with other native TFs. The effect of ARE arrays was most prominent in longer ARE array plasmids containing more than 64 repeats. Moreover, we observed an inverse relationship between NRF2 and NF-κB in the presence of ARE arrays. Our work illustrates examples of ways that TF RE arrays can be used as tools to not only engineer TF responses but also observe TF interactions.

While targeting NRF2 required longer ARE arrays, synthetic TF rtTA significantly perturbed expression with just short TetO arrays. Similarly, shorter arrays of lentiviral integrated Prrx1 and Tead1 did not achieve the desired effects.

A low number of arrays in cells also reduced observations of gene regulation with native TFs as seen when adding equimolar NRF2×12 array compared to gene of interest. Transiently transfecting four times the NRF2×12 array led to a reduction in gene expression. As such the few, random multi-locus integrations of shorter Prrx1 or Tead1 RE arrays may also have contributed to maintenance of fibroblast identity not being sufficient to elicit the desired regulatory outcomes. Therefore, achieving desired results with RE arrays may require adjusting the number of arrays in cells, RE repeat number, and RE sequence.

This observation has significant implications when using synthetic promoters containing TF REs, which many use to drive gene expression and report on cell states (Johansson et al., 2017; Wu et al., 2019). Careful consideration must be given to the potential impact of additional REs on host cell responses, as they could perturb the system they are built to report on.

The developed CTRL method facilitates the use of RE arrays in various applications. Looking more broadly, repetitive DNA has become of more interest in recent years as researchers try to understand the various roles of repeat sequences in non-coding regions of the genome. Recent studies have outlined how repetitive genomic DNA plays a role in regulating gene expression (Horton et al., 2023; Naqvi et al., 2023; Stankey et al., 2024), bridging DNA structure (Zhang et al., 2023) and providing genetic diversity (Liao et al., 2023). As such, we have shown a first use of RE array plasmids in mammalian cells and the effect of RE arrays on regulatory frameworks, but this only represents one use, and more uses of CTRL and RE arrays are possible. For example, RE arrays may provide tools to engineer and understand diseases related to repetitive elements, to control cellular programming by altering TF availability, to research the effects of DNA structures in cells and to map TF relationships.

## Materials & Methods

### Cell Culture

HEK-293T (ATCC) cells were cultured in DMEM (Gibco, 31966047) supplemented with 10% FBS (Gibco, 16000-044). CHO-K1 cells were cultured in α-MEM (Sigma Aldrich, M4526) supplemented with 10% FBS, 1% Non-essential amino acids and 1% L-Glutamine. All cells were kept at 37°C, 5% CO_2_ in a humidified incubator.

### Molecular Cloning

#### Building Expression Cassettes

To generate pTet-I promoter, PCR was used to add BsmbI sites for seamless cloning of 2 TetO sites into CMV-mKate (Di Blasi et al., 2023). The operator sites were generated by synthesis and annealing of oligonucleotides (IDT) (FW: gagctccctatcagtgatagagatctccctatcagtgatagaga, RV: accatctctatcactgatagggagatctctatcactgataggga) and overhangs compatible and predicted high efficiency ligation (Potapov et al., 2018). Briefly 1 µg of oligos were mixed with T4 ligase buffer in a 50 µL reaction and heated to 95°C for 2 mins and slowly cooled to 25°C, over 50 mins. Vector backbone was amplified to introduce BsmbI sites using primers (AACGTCTCatggtttagtgaaccgtcagatcgaattc, AACGTCTCagctctgcttatatagacctcccacc) and Phusion polymerase in GC buffer using standard manufactures protocol (NEB, M0530). The purified amplicon was ligated to annealed oligonucleotides at a 1:7 ratio using T4 ligase and transformed into DH5α chemically competent *E. coli*.

Guide RNA targeting TetO was built into the UNISAM vector (addgene plasmid #99866) using available BbsI sites and annealed oligos (caccTACGTTCTCTATCACTGATA, aaacTATCAGTGATAGAGAACGTA) following similar method to above.

#### CTRL Assembly

TF binding array plasmids were built using the developed CTRL method. Products were frequently digested to confirm intact repetitive portion. All DNA assemblies performed were confirmed frequently by DNA sequencing. CTRL backbones were built from pFF145 cloning vector adding flanking restriction sites and gene for the strong expression of RFP in bacteria (sequences in supplementary information).

#### Biobrick

500 ng of insert plasmid was digested in a reaction containing 10 units of PstI-HF (R3140, NEB) and 10 units of XbaI (R0145, NEB). Another 500 ng of insert plasmid was digested in a separate reaction containing 10 units of SpeI-HF (R3133, NEB) and 10 units of EcoRI-HF (R3101, NEB). The same amount of vector was digested with EcoRI-HF and PstI-HF. All restriction digestions were performed in 1x CutSmart buffer (NEB) and incubated at 37°C for 2 hours. Inserts and backbone fragments were purified by agarose gel electrophoresis followed by gel extraction of excised DNA. To reduce background, the backbone was dephosphorylated using QuickCIP (M0525, NEB). A molar ratio of 2:1 insert to vector DNA was used in the ligation reaction, which was carried out in 1x T4 DNA Ligation buffer (NEB) with 200 units of T4 DNA ligase (M0202, NEB). Ligation reactions were incubated at room temperature for 4 hours before transformation into chemically competent DH5α bacteria. Transformants were then plated onto selective LB agar plates.

#### Golden Gate

To assemble using Golden Gate (GG), 15 fmol of each repeat-containing part in positions 1 through 4 were combined with 15 fmol of the receiver vector containing a drop-out cassette and the other antibiotic resistance gene (ie: Kanamycin receiver if repeat inserts were in vectors containing ampicillin resistance). This mixture was combined with 5 units of Esp3I (R0734, NEB) and 100 units of T4 DNA ligase in 1x T4 ligation buffer, to a final volume of 10 µL. The cycling conditions were as follows: incubation at 37°C for 5 minutes, followed by 20 cycles of 37°C for 5 minutes and 16°C for 10 minutes. To ensure maximal ligation, a final incubation at 16°C for 20 minutes was performed. Remaining unassembled sites were digested by incubating at 37°C for 30 minutes, and enzymes were denatured by incubating at 75°C for 10 minutes.

### Plasmid Transfection

HEK-293T (1×10^5^ cells/well) were seeded in 24 well plates in 500 µL of complete media, 24 hours prior to transfection. On the day of transfection 0.4 µg total DNA is diluted in 50 µL OptiMEM per well and 1.2 µL of Xtreme HP Gene reagent added (Merck, 6366236001) before incubation at room temperature for 30 minutes before addition dropwise to wells.

CHO-K1 (8 x10^4^ cells/well) were seeded in 24 well plates in 500 µL of complete media, 24 hours prior to transfection. On the day of transfection 0.5 µg total DNA is diluted in 50 µL of OptiMEM per well and 1.5 µL TransIT-X2 transfection reagent (Mirus, MIR 6003). Further details about transfection conditions are provided in the Supplementary Files.

### Flow Cytometry & Statistical Analyses

Transfected cells were detached from 24 well plates 48 hours post-transfection. Cells were washed and centrifuged before resuspension in DPBS. For Live/Dead staining (Thermo Fisher Scientific, L3224) 0.05 µM calcein AM and 4 µM ethidium homodimer-1 were added and incubated in the dark before flow cytometry measurement.

Cells were added to filter-capped flow cytometry tubes to disrupt clumps and measure using Attune NxT flow cytometer (Thermo Fisher). Cells not transfected with DNA are used to gate singlet population which is applied to all samples. Then for each sample, 20,000 events from this gate are recorded. The geometric mean of fluorescence was calculated using FlowJo and statistical significance of at least 3 biological replicates was assessed using ANOVA tests using GraphPad Prism software.

### Viral transduction and MEF to iMN reprogramming

Retrovirus for the reprogramming cocktail containing Ngn2-Isl1-Lhx3 and HRAS^G12V^-IRES-p53DD was produced in Plat-E cells. Viral supernatant was collected, filtered, and used for transduction of MEFs two days in a row.

Lentivirus was produced using HEK-293Ts. Cells were transfected with packaging, envelope, and transfer plasmids at a mass ratio of 1:2:1. 6-8 hours after transfection, the media was replaced with fresh HEPES buffered media. Lentiviral supernatant was collected twice at 24 and 48 hours after the first media change. After the second collection, the viral supernatant was filtered and was mixed with Lenti-X concentrator and incubated at 4°C overnight. The virus was pelleted by centrifugation at 1,500 x g for 45 minutes at 4°C. The supernatant was removed, and the pellets were resuspended in media to a final volume of 33 μL per 6-well of virus. The virus was used for transduction of MEFs on the same day.

Primary MEFs were seeded at 10k per 96-well onto gelatin coated plates a day before transduction. MEFs were transduced two days in a row with 11 μL of each PlatE retrovirus per 96-well. On the second day MEFs were also transduced with 2 µL of concentrated lentivirus per 96-well. Fresh media was included to reach a final volume of 100 μL per 96-well, and polybrene was added to increase transduction efficiency. One the second day of transduction the plates were also centrifuged at 1500 x g for 90 minutes at 32°C to further increase lentiviral transduction efficiency. At 1 dpi the transduction media was replaced with fresh DMEM + 10% FBS. At 3 dpi, the media was replaced with N3 media (DMEM/F-12 containing N2, B27, Glutamax, BDNF, GDNF, CNTF, FGF, and RepSox). At 14 dpi, the cells were dissociated using DNase and papain and analyzed via flow cytometry. Singlets were gated using FlowJo and exported. Quantification and statistical analysis were performed using Python. Plots were generated using pandas, matplotlib, and seaborn packages, and statistical analysis was done using numpy and statannotations packages.

## Acknowledgements

The authors would like to acknowledge the support of the EPSRC Centre for Doctoral Training in BioDesign Engineering (EP/S022856/1) (to AZ, FC and TE) and the Royal Society Research Grants 2019 Round 1 to FC. Research reported in this manuscript was supported by the National Institute of General Medical Sciences of the National Institutes of Health under award number R35-GM143033 and sponsored by the U.S. Army Research Office and accomplished under cooperative agreement W911NF-19-2-0026 for the Institute for Collaborative Biotechnologies (KEG). BL is supported by the National Science Foundation Graduate Research Fellowship Program under grant No. 1745302.

## Supplementary Information

**Supplementary Figure 1:**
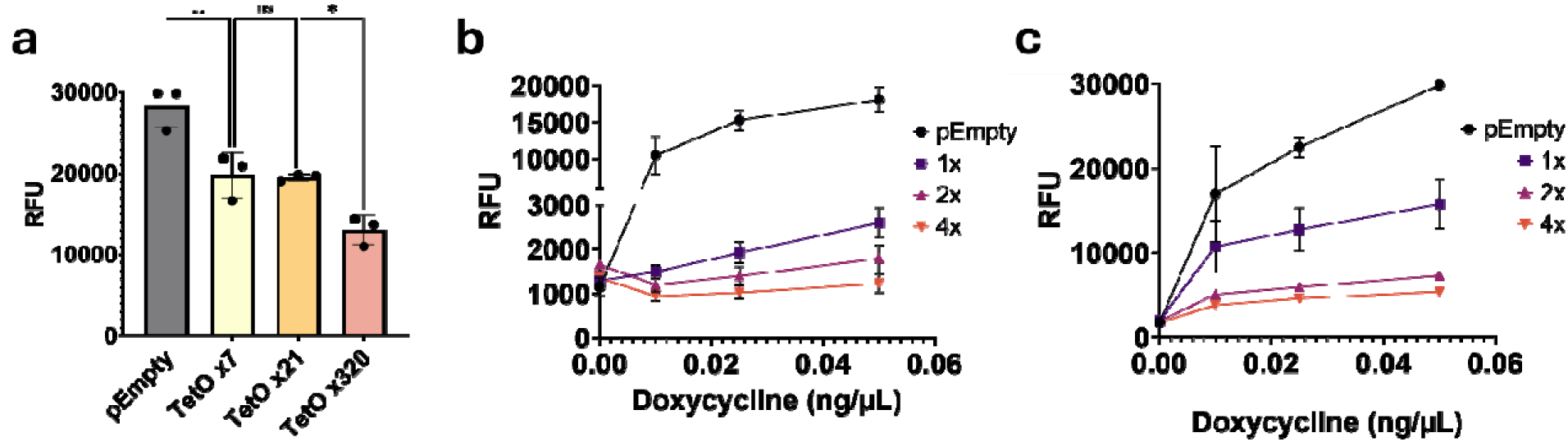
Effect of TetO RE arrays on synthetic circuits in mammalian cells. **a.** In CHO-K1 cells, expression of circuit with EGFP output in the presence of no (pEmpty), 7,21 or 320 TetO arrays in the presence of 0.05 ng/µL doxycycline inducer. **b.** Output expression of EGFP (RFU) in HEK-293T when in the presence of various doxycycline concentrations for no REs (pEmpty), 1x, 2x or 4x TetOx320 array with respect to circuit. **c.** Output expression of EGFP (RFU) in CHO-K1 when in the presence of various doxycycline concentrations for no REs (pEmpty), 1x, 2x or 4x TetOx320 array with respect to circuit. Mean fluorescence of three biological replicates and associated standard deviations are displayed.

**Supplementary Figure 2:**
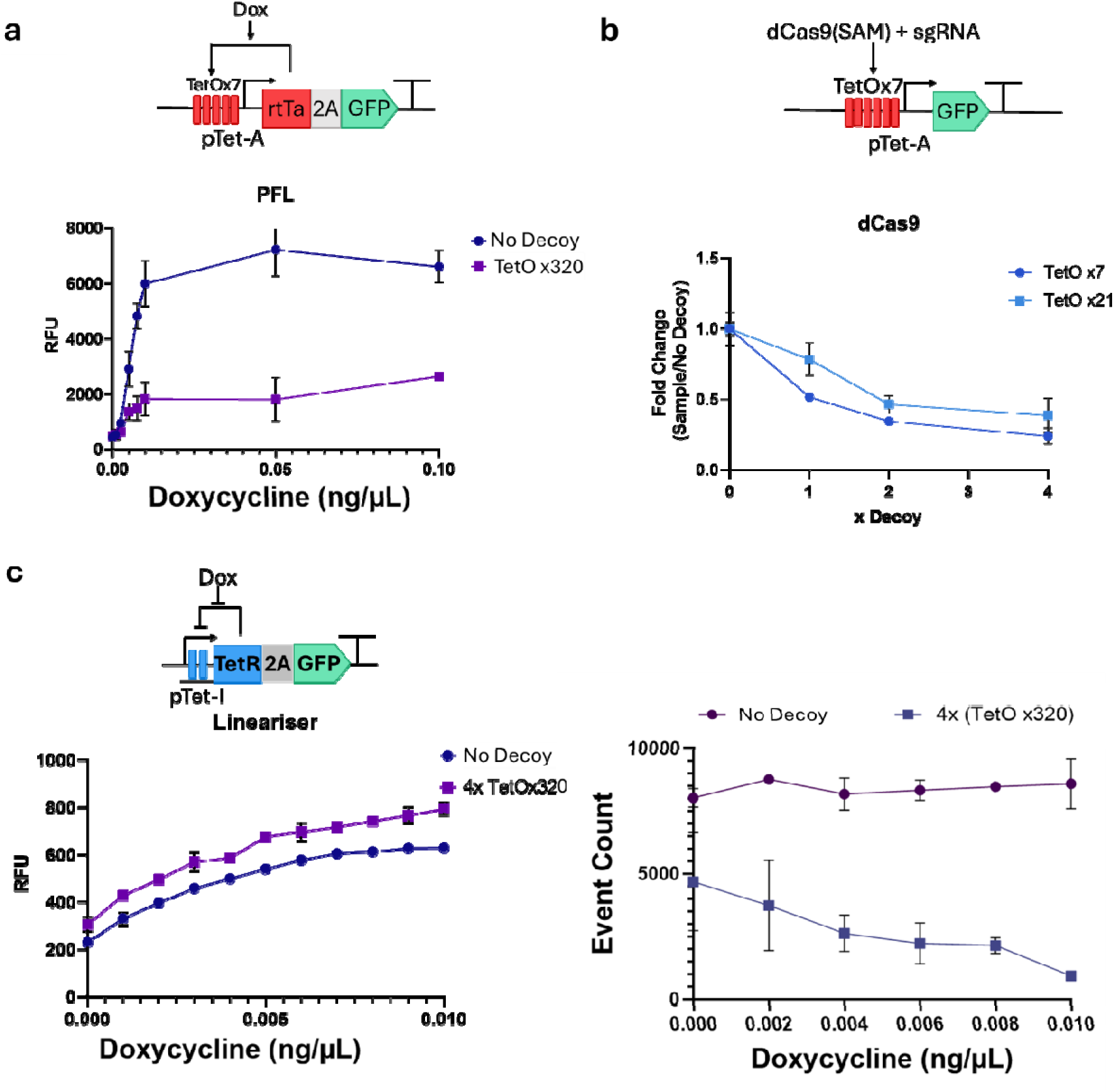
Effect of TetO RE arrays on different synthetic TFs and circuits. **a.** Schematic of positive feedback loop described in Di Blasi et al., 2023 and effect in the presence (purple line) or absence of TetO arrays on reporter output. **b.** Schematic of expression of GFP reporter when dCas9(SAM) and sgRNA targeting TetO sites of the circuit. Graph displaying effects on the fold change of GFP expression in the presence of 1x 2x or 4x arrays containing 7 (dark blue line) or 21 (light blue line) TetO sites. **c.** Depiction of a negative feedback loop described by Szenk et al., in 2020 and effect in the presence (purple line) or absence (blue line) of TetO array sites on reporter output (left) or event count (right). Mean fluorescence of three biological replicates and associated standard deviations are displayed.

**Supplementary Figure 3:**
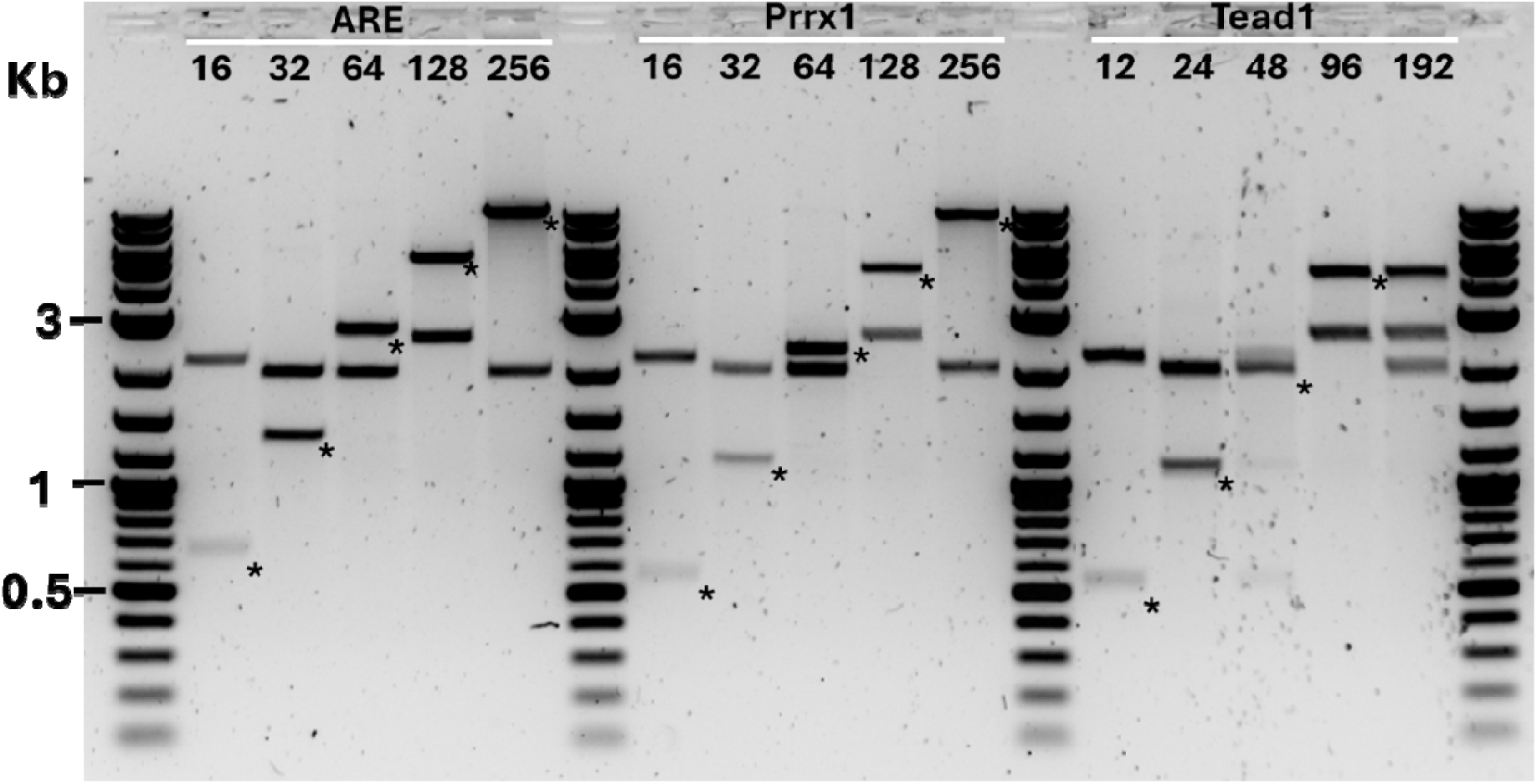
Verification of CTRL assembly of 3 mammalian TF REs. Agarose gel image of EcoRI and PstI restriction enzyme digestion depicting increasing fragment sizes of repeats for three mammalian TFs NRF2, Prrx1 and Tead1 against 1Kb Plus marker (NEB). Repeat containing fragments are marked with an asterisk (*), while the other bands represent either ampicillin or kanamycin resistance containing CTRL backbones.

**Supplementary Figure 4:**
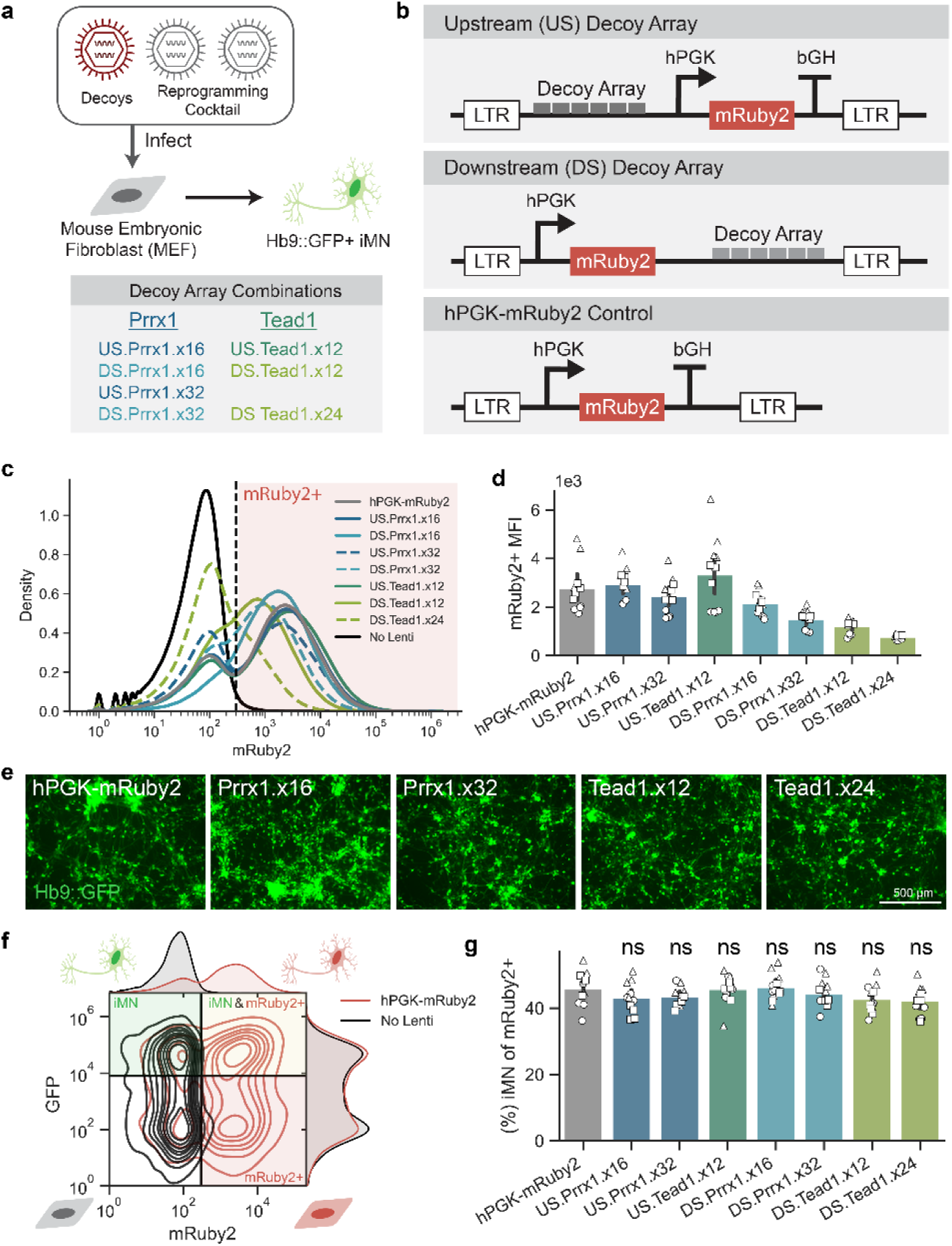
Application of Prrx1 and Tead1 RE arrays. **a**. Schematic of experimental setup. Primary mouse embryonic fibroblasts are transduced with retroviral reprogramming cocktail and lentiviral decoy arrays. Cells express Hb9::GFP when reprogrammed into motor neurons (iMNs). Hb9::GFP and mRuby2 are measured at 14 days post infection (dpi) using flow cytometry. **b**. Schematic of lentiviral vector design including constitutive infection marker and decoy arrays for Prrx1 or Tead1 transcription factors. **c**. Histogram of mRuby2 expression for each vector design. **d**. Mean fluorescent intensity (MFI) of mRuby2+ cells. Mean is shown with 95% confidence interval; marker styles denote biological replicates; n = 3 independent bio reps per condition. **e**. Representative images of Hb9::GFP+ iMNs at 14dpi. **f**. Joint distribution of mRuby2 and Hb9::GFP showing gates for mRuby2+ and iMNs. **g**. iMN purity in mRuby2+ cells at 14dpi. Mean is shown with 95% confidence interval; marker styles denote biological replicates; n = 3 independent bio reps per condition; two-tailed t-test for independent samples, each condition compared to control hPGK-mRuby2, p > 0.05 (ns).

**Supplementary Table 1:**
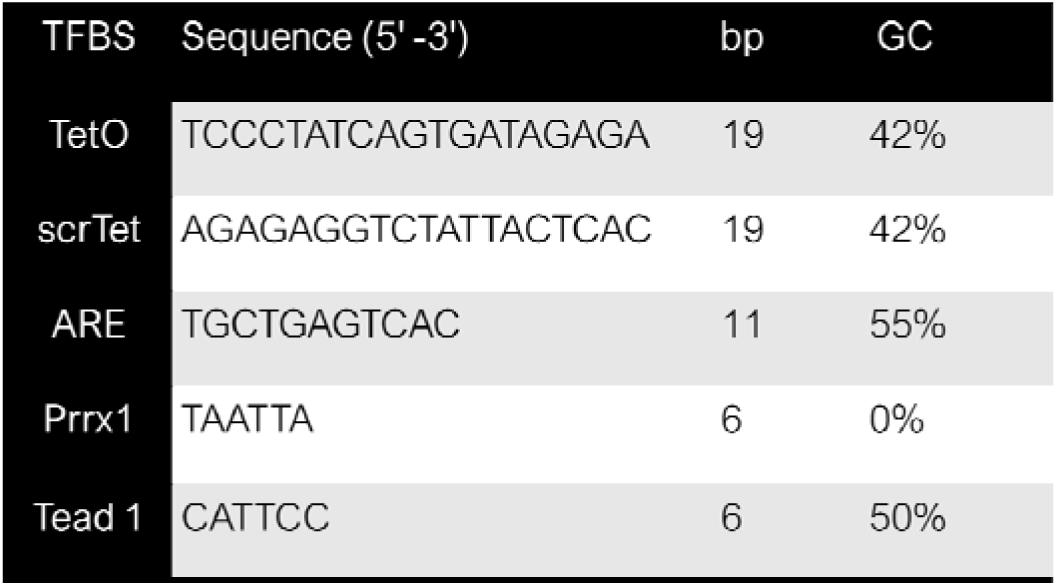
TF DNA recognition elements sequences and properties used for repetitive assembly using CTRL.

**Supplementary Table 2:**
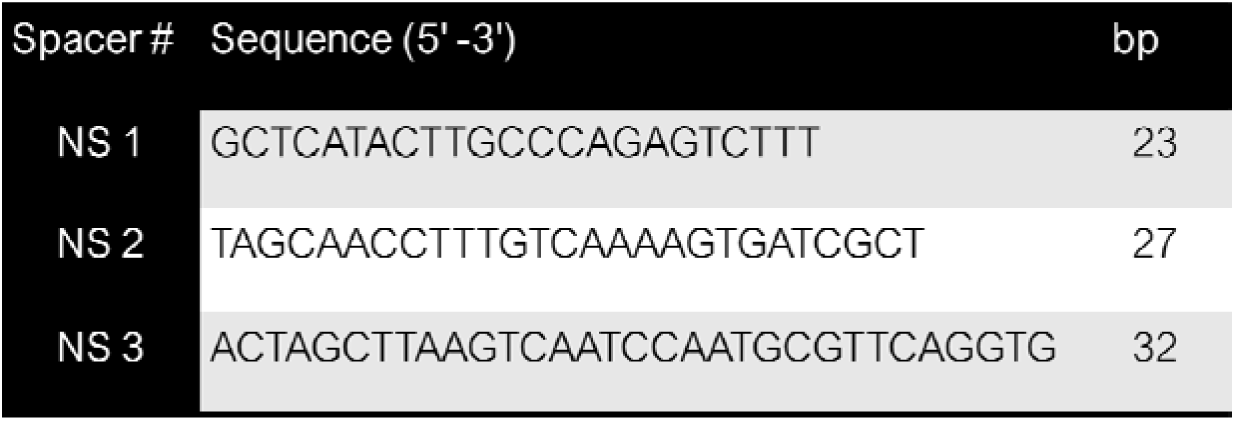
Neutral space sequences used in CTRL, originally described in Wan et al 2020.

**Supplementary Table 3:**
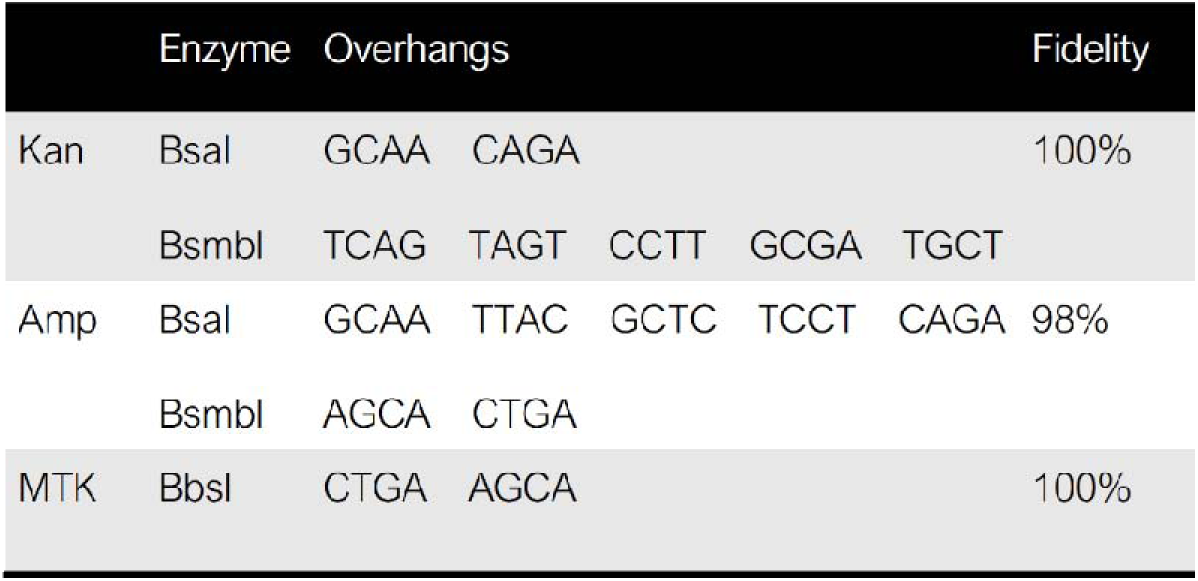
CTRL backbone restriction sites and overhangs and predicted fidelity ligation of these 4bp overhangs.

|  | Enzyme | Overhangs |  |  |  |  | Fidelity |
| --- | --- | --- | --- | --- | --- | --- | --- |
| Kan | Bsal | GCAA | CAGA |  |  |  | 100% |
|  | BsmbI | TCAG | TAGT | CCTT | GCGA | TGCT |  |
| Amp | Bsal | GCAA | TTAC | GCTC | TCCT | CAGA | 98% |
|  | BsmbI | AGCA | CTGA |  |  |  |  |
| MTK | BbsI | CTGA | AGCA |  |  |  | 100% |

